# Investigating the genetic control of plant development under speed breeding conditions

**DOI:** 10.1101/2023.09.04.555916

**Authors:** Nicola Rossi, Wayne Powell, Ian Mackay, Lee Hickey, Andreas Maurer, Klaus Pillen, Karen Halliday, Rajiv Sharma

## Abstract

Speed breeding is a powerful tool to accelerate breeding and research programmes by shortening generation time and has been widely adopted for a range of crop species. Despite its success and growing popularity with breeders the genetic basis of plant development under speed breeding remains unknown. In this study, we explored how genotypes respond in terms of developmental advancement under different photoperiod regimes in the context of speed breeding. A subset of the barley HEB-25 Nested Association Mapping population was evaluated for days to heading and maturity under two contrasting photoperiod conditions: 1) Speed Breeding (SB) consisting of 22 hours of light and 2 hours of darkness), and 2) Normal Breeding (NB) consisting of 16 hours of light and 8 hours of darkness. GWAS revealed that developmental responses under both conditions were largely controlled by two loci: *PPDH-1* and *ELF3*. Allelic variants at these genes determine whether plants display early flowering and maturity under both NB and SB. At key QTL regions, domesticated alleles were associated with late flowering and maturity in NB and early flowering and maturity in SB, whereas wild alleles were associated with early flowering under both conditions. We hypothesise that this may be related to the dark dependent repression of *PPD-H1* by *ELF3* which might be more prominent in NB conditions. Furthermore, by comparing development under two contrasting photoperiod regimes, we were able to derive an estimate of plasticity for the two traits. Interestingly, plasticity in development was largely attributed to allelic variation at ELF3. Our results have important implications for our understanding and optimisation of speed breeding protocols particularly when incorporating genetics from wild relatives into breeding programmes and the design of breeding programmes to support the delivery of climate resilient crops.

## Introduction

The world’s food demand is expected to rise significantly by 2050, by as much as 56% (van Dijk et al., 2021). This increase is primarily due to the combined effects of climate change, population growth, and global food supply disruptions. To meet this demand, it is essential to increase crop yields sustainably (Smith, 2013). This has been achieved in the past through a combination of improved management practices and the generation of superior germplasm (Cooper et al., 2020; Fradgley et al., 2023). However, recent advances in high-throughput genotyping and phenotyping technologies have opened up new opportunities to accelerate the rate of genetic gain in crops (Li et al., 2018). By reducing crop generation time and using marker-assisted selection and genomic prediction, breeders can now significantly increase the yield potential of crops (Gosal et al., 2020). This has prompted breeders to accelerate the seed to seed time of crops through the deployment of technologies to support rapid generation cycling, such as shuttle breeding, single seed descent, double haploid and, more recently, speed breeding (Watson et al., 2018).

Prior to speed breeding, the main technology for faster breeding cycles was the use of double haploid technology (DH), which quickly generated homozygous lines after F1 or F2 generations. However, DH has two main drawbacks: firstly it requires expensive tissue culture labs, secondly DH populations derive from low recombination events and cross-over rates, which increases the population size needed (Inagaki et al., 1998). Moreover, its effectiveness varies among genotypes as often many genotypes are non-responsive to tissue culture (Hooghvorst et al., 2021). Alternatively, single seed descent (SSD) method was adopted in many crops. Traditionally, the SSD approach involves advancing each F2 individual through selfing in a controlled environment with a 16-hour photoperiod for long day plants, achieving up to three generations per year. This method comes with reduced costs and higher genetic variability compared to DH breeding. The increased genetic diversity that results from SSD contributes to improved selection efficiency and serves as a protective measure against genetic drift. As a result, SSD is a powerful tool for enhancing the overall efficacy and success of crop improvement programs. However, SSD is not always a superior alternative to DH as it leads to slower development of recombinant inbred lines (Caligari et al., 1987; Powell et al., 1986).

Although research on the effects of extended photoperiod on plant growth and development began almost a century ago (Arthur et al., 1930; Garner and Allard 1927), it was not until recently that researchers began to investigate the most efficient combination of environmental factors for reducing the breeding cycle. Watson et al., (2018) demonstrated that speed breeding methods could be adapted to reduce generation time for a broad range of crop species. Developing an efficient speed breeding protocol involves optimising several environmental factors, with a key one being the exposure to prolonged photoperiods for long-day species. Speed breeding can be integrated with other technologies to achieve different breeding objectives such marker assisted selection (MAS) for simple traits as genomic selection (GS) for complex traits (L. T. Hickey et al., 2017, 2019; Pandey et al., 2022). A body of research has advanced speed breeding protocols aiming to reduce the breeding cycle in long- and short-day plants using controlled environments (Cazzola et al., 2020; Chiurugwi et al., 2019; Fang et al., 2021; Watson et al., 2018; J. M. Hickey et al., 2017; Mobini et al., 2020; Samineni et al., 2020; Schilling et al., 2023). Depending on the species, an optimized protocol can reduce the time from crossing to testing to 18 months or two years, much shorter than SSD or shuttle breeding. Furthermore, speed breeding offers significant advantages over DH technology as it maintains higher recombination and cross-over events while still achieving a similar reduction in generation time at a reduced cost. As a result, rapid cycling protocols have become popular in plant breeding programs around the world.

Despite the recent success of speed breeding, there are still opportunities for refinement by optimizing energy and management costs, as the tool is still in its infancy. This is intertwined to the limited understanding underlying the genetics of plant development under such conditions. Specifically, we do not know whether flowering and maturity under very long days (e.g., 22 hours light in speed breeding conditions) is genotype-dependent and under different genetic controls compared to standard long days (e.g., 16 hour days). Understanding this could help breeders and researchers to develop more effective speed breeding protocols. Enhancing our knowledge on this matter can significantly influence the decision-making process for breeders and researchers when considering the adoption of this technology, leading to more effective and targeted crop improvement and research strategies. While studies on speed breeding in cereals have shown that plant development can be accelerated under these conditions (Cha et al., 2022; Watson et al., 2018), experiments have mainly focussed on modern or elite germplasm. As introgression breeding is becoming a valuable tool for gaining access to wild genetic diversity that can help crops adapt to climate change (Gramazio et al., 2021; Hao et al., 2020; Hernandez et al., 2020; Khan et al., 2023; Zhang et al., 2023). Therefore, a better understanding of the genetics of speed breeding would help pre-breeders develop protocols that are effective in these programs.

To shed light on the genetic basis of speed breeding, the present study examined the ‘Halle Wild Barley’ (HEB-25) nested associated mapping (NAM) population, which segregates for both wild and domestic alleles (Maurer et al., 2015). The lines were phenotyped for key developmental traits under both **speed breeding** (22 hours of light and 2 hours of darkness) and **normal breeding** (16 hours of light and 8 hours of darkness). Data from these experiments and whole-genome marker data using the Infinium iSelect 50k SNP chip (Maurer and Pillen 2019) was used in Genome Wide Association (GWAS) to identify genetic loci associated with the differential responses of spring barley lines grown under the two different artificial growth conditions. To our knowledge, this is the first study to identify the genetic basis of plant development under speed breeding, providing insight into the mechanisms controlling the plant’s development-related traits under long and very long days. The results of this study have important implications for the deployment of speed breeding to accelerate the utilisation of genetic diversity, particularly wild relatives, to support the development of future crops.

## Material and methods

### Plant material

The present study uses the multiparent nested associated mapping (NAM) population ‘Halle Wild Barley’ (HEB-25), developed by Maurer et al., (2015). This population was generated using 25 wild barley parents (24 *Horderum vulgare ssp. spontaneum*, Hsp and 1 *Hourderum vulgare ssp. agriocrithon*) crossed with spring barley cultivar Barke (*H. vulgare ssp. vulgare*, Hv). The resulting generation was backcrossed with the female parent Barke, following three generations of selfing through single seed descent (BC_1_S_3_). Thereupon, the deriving lines were propagated through the 6^th^ generation of selfing (BC_1_S_3:6_). Further details on the population development is provided in Maurer et al., (2015). This multiparent NAM population has become a crucial genetic resource for investigating various essential traits in barley, including stress tolerance and yield (Büttner et al., 2020; Mehnaz et al., 2021; Saade et al., 2016; Sharma et al., 2018; Wiegmann et al., 2019). In our study, we aimed to efficiently evaluate the HEB-25 to study the genetics of speed breeding. However, screening the entire population in a glasshouse posed practical limitations. To overcome this issue, we implemented a random sampling approach to select a subset of 190 genotypes from the population, consisting of three to four genotypes from each of the 25 families present in the population. To select a subset of 190 genotypes from the HEB-25, we employed the RAND() function available in Microsoft Excel version 2010 (MS-Office). The use of this function allowed to randomly select three to four lines from each of the 25 families, thus minimizing the potential for bias in our selection process. To ensure that we selected an extensively varied subset, we conducted a principal component analysis (PCA) with R studio version 4.2.2. This analysis employed the complete panel along with an SNP matrix consisting of 32,955 markers. Subsequently, the PCA plot was produced using the R-package “ggplot2” (Wichham 2016) As depicted in Figure S1, our dataset comprehensively represents the entire population and exhibits considerable diversity. The selected subset was screened in two subsequent experiment rounds (1^st^ from November 2021 to March 2022, 2^nd^ from July to October 2022). A set of 12 genotypes were included in both rounds of screening for normalization of the experiments that was subsequently used to combine the data across the two screening rounds via a linear mixed model, as outlined in the “statistical analysis” section.

### Speed breeding experiments & phenotyping

In order to fulfil the aim of this study, we gathered phenotypic data on the development of barley plants under different controlled environmental conditions. To achieve this, the experiments were conducted in a glasshouse located at SRUC’s Peter-Wilson campus (55°55’17.386” N −3°10’42.175” E) manufactured by CambridgeHOK. By measuring developmental traits of the plants under both conditions, we aimed to gain insights into the genetic characteristics of plant development under speed breeding.

The experimental conditions were meticulously chosen to ensure that the phenotypic data collected accurately represented the impact of the prolonged photoperiod used in speed breeding for cereals. The first glasshouse compartment had a photoperiod of 16 hours of light and 8 hours of darkness (16:8) (hereon called **Normal Breeding**: **NB**), while the second compartment was set up for speed breeding and had a photoperiod of 22 hours of light and 2 hours of darkness (22:2) (hereon called **Speed Breeding**: **SB)**. The temperature in both compartments was programmed at 22 degrees Celsius during the day and 17 degrees Celsius at night, in accordance with the specifications of Watson et al., (2018). The experimental unit was one plant per 0.3 litres pot at a density of sowing of 77 plants/m^2^, with five replicates per genotype in a complete randomized block design (RBD). The plants were distributed across the benches in 50 columns and 10 rows of each treatment. The glasshouse is supplied with 400W High Pressure Sodium light fixtures (Sylvania GroLux). The light intensity and the temperature were measured via a quantum sensor (SKP 200 – Skye Instruments) and dataloggers (EasyLog USB), respectively.

A set of 100 HEB lines were sown in November 2021, and another set of 96 HEB lines in July 2022. Twelve HEB lines were cultivated under both experiment rounds for normalization, as detailed in the statistical analysis section.

Our study concentrated on two traits that have high heritability and are essential for the successful completion of barley’s life cycle and development: days to heading (as a proxy for flowering time) and days to maturity. We scored the traits by measuring the number of days it took for the plant to reach growth stages BBCH49 (Heading - HEA) and BBCH92 (Maturity – MAT) using the BBCH scale developed by Lancashire et al., (1991) under both NB and SB conditions. Additionally, we measured phenotypic plasticity, which is defined as the changes exhibited by a genotype when grown in different environments (Laitinen et al., 2019). Hence, in our study, plasticity is the quantification of changes in developmental advancement of a genotype across the two controlled environment conditions. Plasticity was calculated for each genotype as the difference between the average trait performances under NB and SB. We utilized these derived traits to identify genetic factors that contribute to the plasticity of HEA and MAT (Plasticity.HEA and Plasticity.MAT) across NB and SB. This is a useful measure of adaptation, particularly in light of the changing global environment characterized by abiotic stresses. Gaining insights into the genetic basis of differential responses observed in long days and very long days can help us understand how plant development varies under different light conditions. This understanding can be used to develop speed breeding protocols that are tailored to specific genetic backgrounds or germplasm pools (e.g., elite versus wild).

### Statistical analyses

After checking the phenotypic trait values manually for typographical errors, we excluded outliers exceeding 3 standard deviations in each genotype. Subsequently we removed genotypes with less than 3 replicates, per environment, from the analysis. Next, we fitted the best linear mixed model to obtain the Best Linear Unbiased Estimator (BLUE) for the studied traits, considering genotypes as fixed effects and the experiment round, along with the row and column effects (due to the varying light distribution across benches), as random effect. Cultivating 12 common genotypes across experiment rounds and incorporating this factor into the model enabled the normalization of phenotypic data from both rounds of the experiment. The models were fitted using “lmer” function from the package “lme4” (Bates et al., 2015) in Rstudio version 4.2.2. We then compared different models that considered either row and/or column effects or none of them and selected the best performing model based on the lower AIC (Akaike Information Criterion). The model comparison was made via the “aic” function in the basic package “stats” in Rstudio version 4.2.2.

Summing the genotypes effects to the intercept provided unique values for each genotype which were then used for the GWAS and for calculating plasticity.

Traits’ heritability was calculated using Piepho’s (Piepho and Möhring 2007) method using the R-scripts provided in Covarrubias-Pazaran (2019).

The GWAS was performed using barley 50K SNP markers (Bayer et al., 2017; Maurer and Pillen 2019) by fitting following model:

Y = Xb + Wm + Zu + e

where **y** is a N × 1 column vector of the BLUE values of phenotypic data of N NAM lines (N = 190 max in our case); b is a vector of population structure effects as fixed effects; X is an incidence matrix relating b to y, consisting of principal components loadings from the PCA; **m** is a vector of fixed marker effects; **W** is a marker matrix containing marker types (as −1, 0 and 1); **u** is a vector of random polygenic effects where 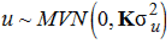, **K** is the additive relationship matrix obtained from the markers using the function “A.mat” in the “rrBLUP” package (Endelman 2011) in Rstudio version 4.2.2: **Z** is an incidence matrix linking **u** to **y**; **e** is a vector of random residuals where 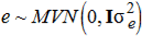 and **I** is the identity matrix.

The correction for population structure was conducted via the kinship correction and using the top 6 principal components as covariates, namely the Q+K model (Isidro-Sánchez et al., 2017). The number of principal components used in the analysis was established from a scree plot and by visually evaluating the component number at which the rate of eigenvalue decrease began to plateau.

GWAS was conducted using SNP with MAF > 0.05 and the threshold of False Discover Rate (type I error rate) was set at α = 0.05 for each trait. More details on the markers 50k Illumina Infinium iSelect SNP array given in (Maurer and Pillen 2019).

In addition, markers effect size was computed using the “mixed.solve” function in “rrBLUP” using Rstudio version version 4.2.2. The effects are derived from the wild parents’ of the population.

### Analysis of alleles associated with *PPD-H1* and *ELF3*

Genotype groups were created from polymorphisms present at some of the significantly associated markers in the two major QTLs (co-located with the candidate genes *ELF3* and *PPD-H1*) found in the GWAS scans. This yielded 4 different groups based on the allelic combinations for the SNPs in the two loci. Being Barke the only domesticated parent for the HEB-25 and used as reference genome for the SNP computation, the alleles presenting polymorphism to this genome are referred as “Hsp” (from *H. sponaneum*, wild parent) and the Barke ones as “Hv” (from *H. vulgarae*, domesticated parent) as in Table 1. These four groups were then displayed via boxplot for all the traits. A pairwise student t-tests was used for detecting differences among the genotypic groups. All the comparisons between the groups were made both in form of parametric t-test and permuted t-test. The boxplots were created using the package “ggplot2” (Wickham 2016) and t-tests via the “t.test” function in Rstudio version 4.2.2.

**Table 1.**
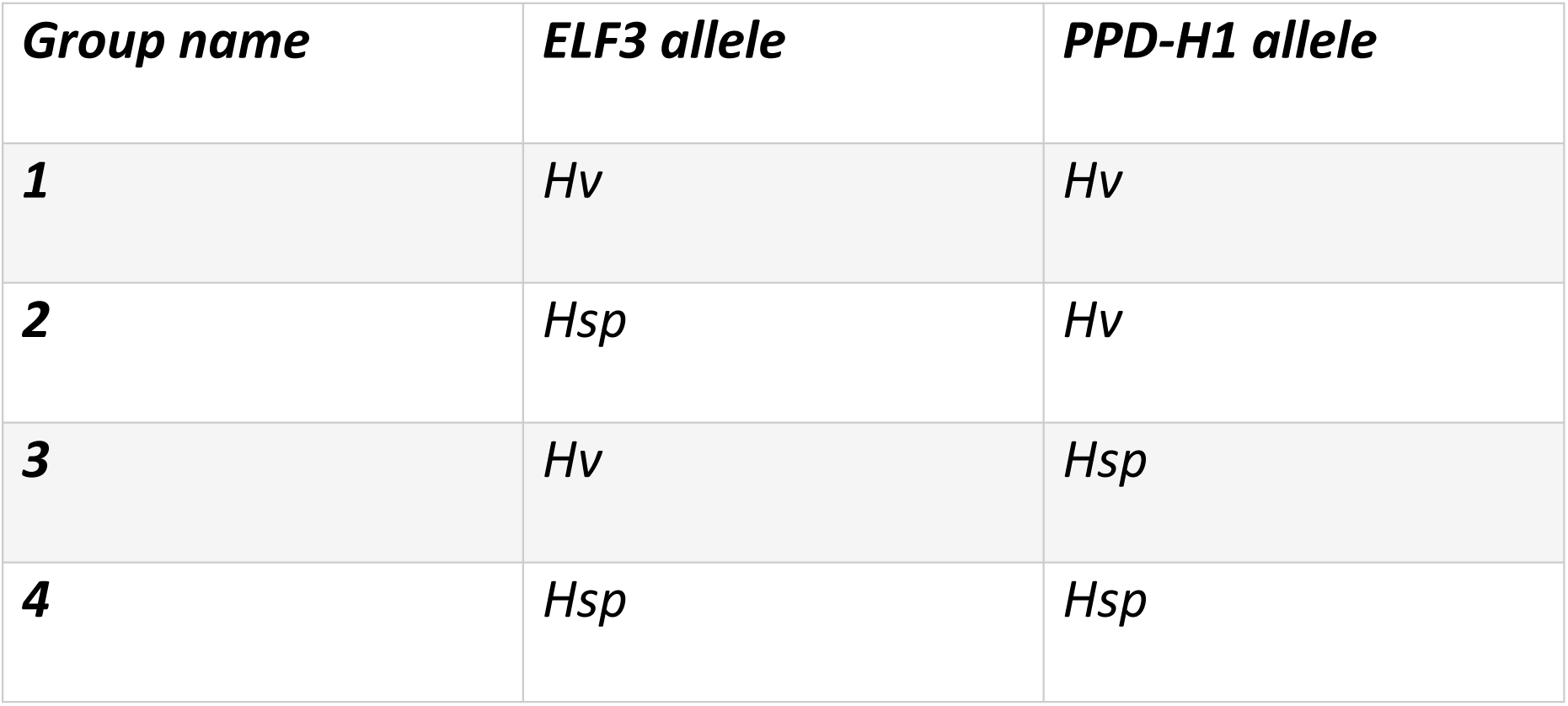
Genotypes groups based on the combinatorial allelic combination from the haplotypic analysis. Hv: domesticated allele; Hsp: wild allele. These haplotypes are derived from SNPs underestimating original haplotypes harboured in the HEB25 families as we do not have access to sequence information yet.

| **Group name** | **ELF3 allele** | **PPD-H1 allele** |
| --- | --- | --- |
| **1** | *Hv* | *Hv* |
| **2** | *Hsp* | *Hv* |
| **3** | *Hv* | *Hsp* |
| **4** | *Hsp* | *Hsp* |

## Results

### Plant Development Acceleration due to Speed Breeding is Genotype Dependent

In general, plants completed their life cycles faster in SB than in NB. Flowering (HEA) occurred 36 ± 7 days after germination in SB conditions and 52 ± 11.5 days in NB conditions. This corresponds to a 15.9 ± 6.88-day developmental acceleration under SB. However, the average difference in days to maturity (MAT) between the two conditions was 7.7 ± 6.88 days. **Figure S2** shows the distribution of these traits as frequency histograms. BLUE values for HEA and MAT and their derived plasticity traits, along with the summary statistics for mean, standard deviation, minimum and maximum values, and heritability values are provided in **Data S1 and Table S1**, respectively.

Notably, a significant proportion of the lines (approximately 90%) flowered and matured earlier under SB than under NB. This suggests that there is substantial variation in trait values and that SB has an important effect on plant growth.

Plants subjected to SB exhibited a faster completion of their life cycles compared to those under NB. This effect was evident under both flowering and maturity stages. Importantly, a substantial amount of genetic variability was observed in how plants responded to both conditions, enabling a GWAS to be conducted as described in the subsequent section. The heritability values of these traits were high, albeit lower in SB than in NB (**Table S1**).

These findings indicate that certain genotypes display early flowering only during prolonged photoperiods, whereas other are early flowering under both long days and very long days, suggesting the involvement of specific genes that regulate plant development.

### The effect of allele combinations associated with *ELF3* and *PPD-H1* on MAT, HEA, and Plasticity

To better understand the genetic factors that control plant development under SB and NB conditions, a GWAS was conducted for each trait. We focused on identifying QTLs associated with the regulation of the development under SB and NB conditions. Specifically, six GWAS scans were performed across four primary traits: HEA in NB, HEA in SB, MAT in NB, MAT in SB and their corresponding plasticity traits: Plasticity.HEA and Plasticity.MAT, across the two conditions. Manhattan plots and the list of markers, their position, the level of association −log_10_(*P-*value) ≥ 4 and their effects are provided in **Figure 1** and **Data S2**, respectively.

**Figure 1.**
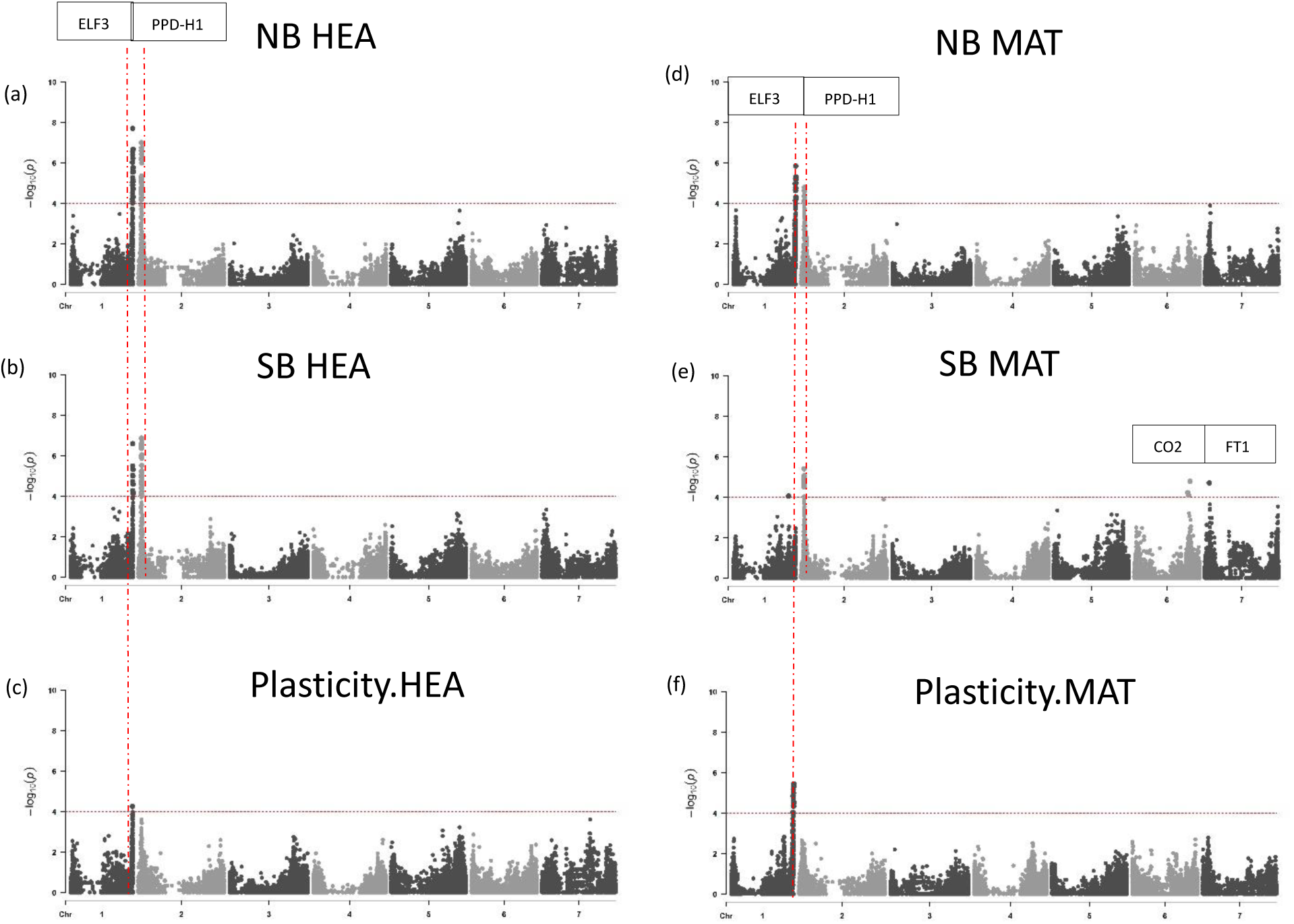
Manhattan plots from the six-traits (a-f). Seven barley chromosomes are shown (1H-7H) horizontally and –log10(p-values) are displayed vertically by dotted line. Significant FDR threshold grey dashed line set at 0.05. The coinciding flowering time candidate genes are shown in the rectangle boxes. Plots were created using the “CMplot” package (Yin et al., 2021) in R studio version 4.2.2. The details of the significant peaks and the markers underlying these peaks are provided in **Data S2**.

The GWAS scans of HEA and MAT traits under both SB and NB revealed two prominent QTLs that are co-located with the major flowering-time genes *ELF3* on chromosome 1H and *PPD-H1* on chromosome 2H (Russell et al., 2016), implicating their central importance in the control of flowering and maturity. Interestingly the *PPD-H1* association with MAT was maintained under both NB and SB. Conversely, for the plasticity trait, the *ELF3* association remained significant, highlighting its involvement in governing the plasticity of HEA and MAT under SB conditions.

Our GWAS results emphasise the importance of major flowering-time genes in barley for the regulation of HEA and MAT traits, which aligns with previous studies (He et al., 2019; Maurer et al., 2015, 2016). However, our findings also highlight the wider relevance of these two genes specifically in the context of speed breeding, which has not been previously reported in the literature. Furthermore, the identification of two additional QTLs associated with the MAT trait in speed breeding, located on chromosomes 7H and 6H in the regions of *FLOWERING LOCUS T FT1* and *CONSTANS 2 CO2*, respectively, indicated the potential involvement of additional genes in the control of speed breeding. The detection of these QTLs in SB is particularly interesting, considering the strong correlation observed between MAT and HEA traits (**Figure S3, Table S2**). The presence of few additional regions of relevance suggests that these specific regions may have a greater influence on the MAT trait under SB, as they do not exhibit significant association in NB. This finding implies the existence a of unique genetic mechanisms that regulate MAT trait responses in the context of speed breeding.

The validity of our findings was further supported through the incorporation of major QTL peaks from chromosome 1H and 2H as covariates in our GWAS model (**Figure S4, Data S5**). As expected, these two major QTLs disappeared after incorporation as covariates. Consequently, we successfully detected a prominent QTL peak proximal to the *FT1* genomic region on chromosome 7H, indicating a significant association with both HEA and MAT traits. Additionally, we observed smaller QTL peaks in other regions such as chromosome 1H (MAT in SB, candidate genes *PPD-H2* and *GA20ox*) and 5H (HEA in NB, candidate gene *CO2*).

The conspicuous association detected near the *FT1* genomic region on chromosome 7H strongly suggests its importance in regulating HEA and MAT traits in addition to *PPD-H1* and *ELF3*. Furthermore, the significant association of markers in regions other than the ones containing the two main QTLs identified, implies the involvement of the additional genes *PPD-H2*, *GA20ox* and *CO2* in the regulation of these traits.

Overall, our GWAS results shed light on the crucial role of major flowering-time genes in controlling HEA and MAT traits in barley. Additionally, they reveal the broader significance of these genes in the specific context of speed breeding, providing valuable insights not previously reported in the literature. The identification of QTLs associated with the MAT trait in SB further suggests the involvement of additional genes, highlighting the complexity of this trait and its response to different growth conditions. In fact, it is important to mention that the effect of hand watering on MAT is more prominent than its effect on HEA (Qaseem et al., 2019).

### Domesticated Alleles at *ELF3* and *PPD-H1* Confer Higher Plasticity

The combination of alleles at the *ELF3* and *PPD-H1* genes are important under both SB and NB conditions. Different allelic combinations at *ELF3* and *PPD-H1* loci (**Table 1**, *PPD-H1Hv/ELF3Hv* - group 1, *PPD-H1Hv/ELF3Hsp* - group 2, *PPD-H1Hsp/ELF3Hv* - group 3, *PPD-H1Hsp/ELF3Hsp* - group 4) also affect MAT, HEA and their plasticity. As observed by Maurer et al., (2015) and Zahn et al., (2023), genotypes carrying at least one wild allele at one of the two loci (groups 2,3 and 4) tend to flower earlier compared with lines carrying domesticated alleles under both loci (group 1, p. values <=0.0009, **Figure 2**). This is consistent with the wild alleles effect of the significant markers found in our GWAS scan, as their effect is negative on the traits value, accelerating the plant’s development (**Data S2**).

**Figure 2.**
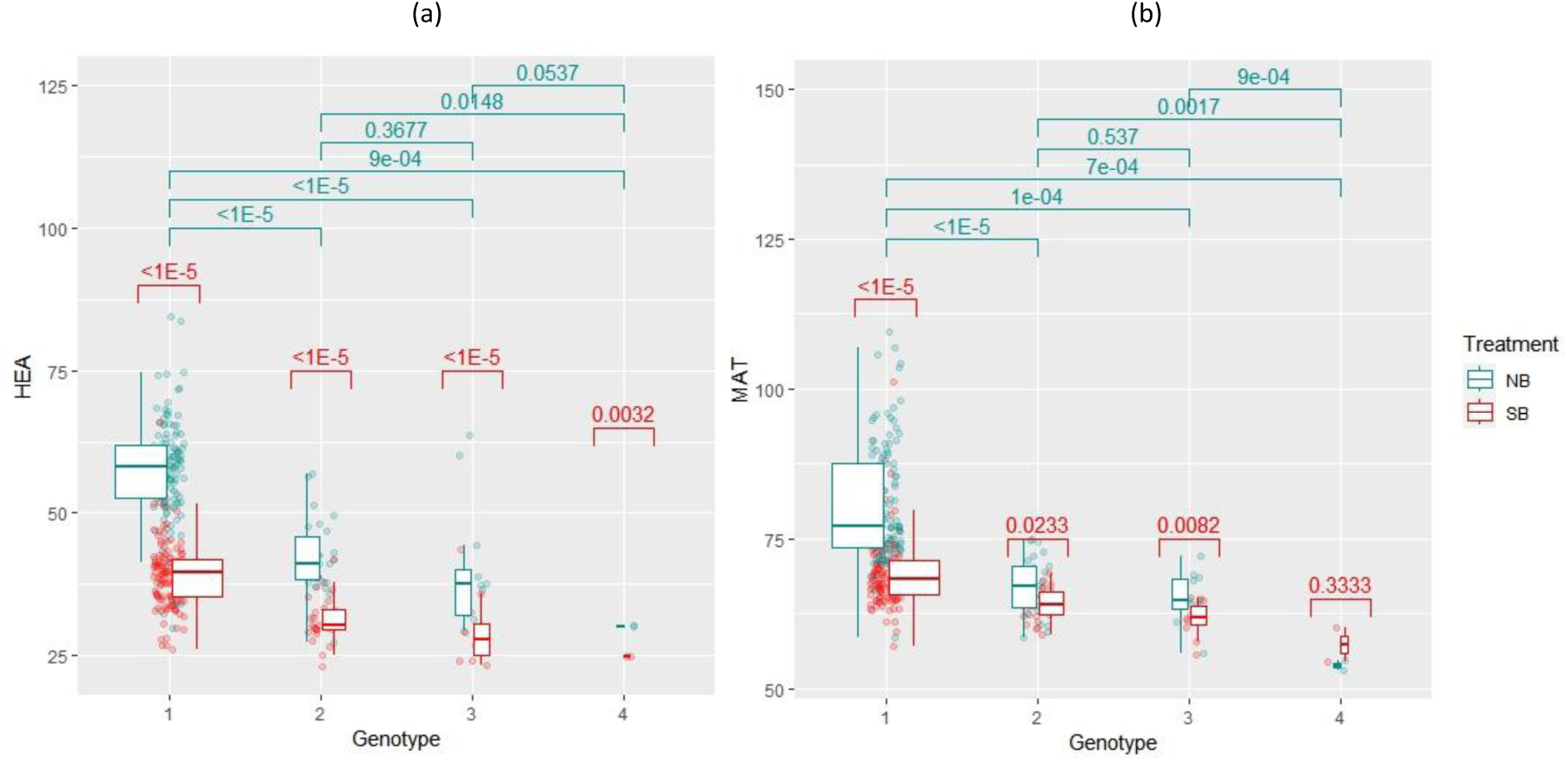
PPD-H1 and ELF3 alleles-based box plots from HEA (a) and MAT (b). Boxplots of the response of different genotype groups to MAT and HEA, arising from the combinatorial allelic analysis. PPD-H1_Hv_/ELF3_Hv_ - group 1, PPD-H1_Hv_/ELF3_Hsp_ - group 2, PPD-H1_Hsp_/ELF3_Hv_ - group 3, PPD-H1_Hsp_/ELF3_Hsp_ - group 4. P-values shown are from permutation t-tests.

Domesticated haplotypes at both loci appear to give higher levels of plasticity compared to the wild haplotypes (p.values<=0.02, **Figure 3, Data S3a**). Nevertheless, the means between experimental conditions within all genotypes’ groups are significantly different except for genotype group 4’s MAT. Therefore, SB reduced the HEA and MAT in all the genotypes studied, however, looking at the means in **Data S3b** and **Figure 3**, the extent of cycling acceleration in genotypes carrying wild alleles is very low compared to the ones harbouring domesticated alleles, especially for MAT. Hence, wild alleles at *PPD-H1* and *ELF3* confer early flowering under both conditions.

**Figure 3.**
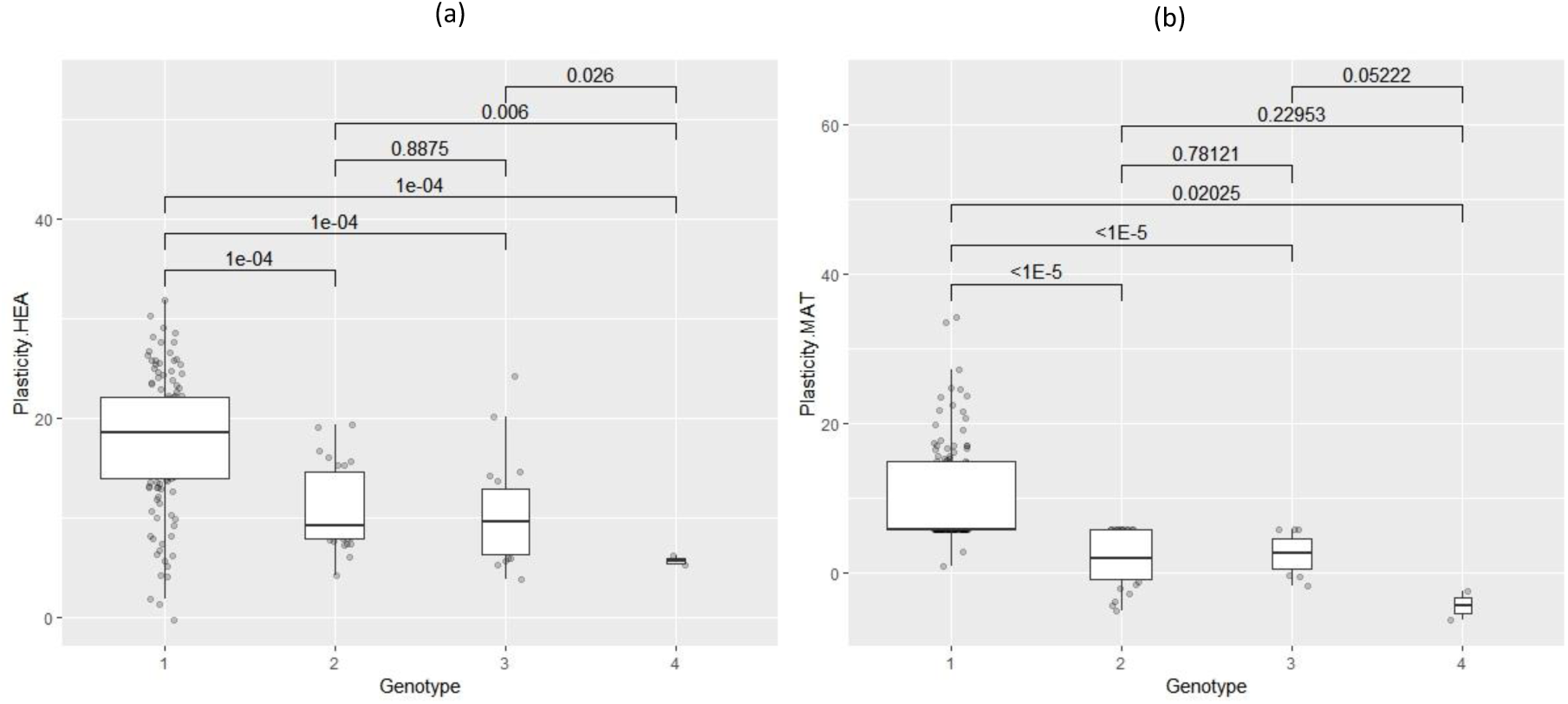
PPD-H1 and ELF3 alleles-based box plots from Plasticity. HEA (a) and Plasticity.MAT (b). Boxplots of the response of different genotype groups to Plasticity.HEA and Plasticity., arising from the combinatorial allelic analysis. PPD-H1_Hv_/ELF3_Hv_ - group 1, PPD-H1_Hv_/ELF3_Hsp_ - group 2, PPD-H1_Hsp_/ELF3_Hv_ - group 3, PPD-H1_Hsp_/ELF3_Hsp_ - group 4. Significance shown are from permutation t-tests.

Finally, we ranked the genotypes based on the MAT value under SB and examined the genotype group to which the latest maturing genotypes belonged. This analysis is valuable in investigating whether breeders tend to unintentionally select for specific alleles at *PPD-H1* and *ELF3* if they do not advance the latest maturing plants group in the population. Later maturing plants in the 75^th^ percentile harboured domesticated alleles at *ELF3* and *PPD-H1* genes (**Data S5, Figure S5**).

## Discussion

To our knowledge this is the first time that the underpinning genetic control of speed breeding has been explored. Various studies have described the benefits of speed breeding (Ahmar et al., 2020; Bhatta et al., 2021; Bohra et al., 2020; Pandey et al., 2022; Samantara et al., 2022; Song et al., 2022; Wanga et al., 2021) and optimised protocols for the deployment of speed breeding (Cazzola et al., 2020; Chiurugwi et al., 2019; Fang et al., 2021; Watson et al., 2018; J. M. Hickey et al., 2017; Mobini et al., 2020; Samineni et al., 2020; Schilling et al., 2023) but the genetic basis of plant development under such conditions remains unexplored. In this study we adopted a GWAS approach to unravel the genetic control of speed breeding in a barley NAM population grown under two growth conditions, one with a photoperiod of 22 hours of light 2 hours of darkness (SB) and the other on 16 hours of light and 8 hours of darkness (NB). By studying a subset of the spring barley HEB 25 NAM population (Maurer et al., 2015) a broad range of wild and domesticated alleles were explored and led to the identification of candidate genes associated with the control of developmental traits under both SB and NB. Two significant candidate genes were pinpointed: *ELF3* and *PPD-H1* controlling both days to heading and days to maturity. Most importantly, by measuring the changes exhibited by an individual genotype over the two treatments we were able to derive an estimate for plasticity and ascribe a candidate gene *ELF3* that is strongly associated with it, supporting its role as a key hub integrating gene networks influencing overall plasticity (Laitinen et al., 2019).

Previous studies have shown that the *PPD-H1* and *ELF3* genes are involved in the genetic control of several agronomic traits in barley (Digel et al., 2016; Ejaz and von Korff 2017; Gol et al., 2021; Ochagavía et al., 2022). At the *PPD-H1* locus, a variety of natural variants have been identified, and categorized into two distinct groups: the sensitive allele, *Ppd-H1* which reduces flowering time during long days and the insensitive variant, *ppd-H1* which delays flowering in long days (Russell et al., 2016; Turner et al., 2005; Fernández-Calleja et al., 2021). The former variant likely represents the ancestral allele, found in HEB-25 and present in winter and Australian barleys (Hu et al., 2023) whereas the *ppd-H1* allele is prevalent in Barke and numerous European and North American spring barley cultivars. With respect to *ELF3*, knowledge and understanding of the phenotypic effects of the allelic series is less established than for *PPD-H1*. Faure et al., (2012) were the first to discover a loss of function allele at this locus and they identified *ELF3* as the candidate gene responsible for the *eam8* mutant originating from the Scandinavian induced mutation experiments performed in the past century (Lundqvist 2009). This allele confers early flowering both in short and long days, compared to the domesticated allele. Such response it is similar to the one observed in our study by wild alleles in the HEB 25 that aligns with the findings from Zahn et al., (2023) and Zhu et al., (2023).

Both *PPD-H1* and *ELF3* are linked to the expression of *FT1* and *GA20ox* genes that are downstream floral integrators that control the flowering response (Boden et al., 2014; Campoli et al., 2012; Cheng et al., 2023; Faure et al., 2012; Turner et al., 2005). *ELF3* delays the flowering response whereas *PPD-H1* accelerates the response in long days. In Arabidopsis, *ELF3* is a repressor of *PRR7* (Dixon et al., 2011; Herrero et al., 2012; Nusinow et al., 2011), which is an homologue of *PPD-H1*. In barley, this interaction has been studied at the transcript level (Faure et al., 2012; Zahn et al., 2023) with both the allele present in *eam8* mutant and the wild *ELF3hsp* correlate with a higher *PPD-H1* expression that leads to early flowering compared to the domesticated *ELF3hv* variant. Furthermore, Müller et al., (2020) hypothesized that, as seen in Arabidopsis, *ELF3* antagonizes the light input in the circadian clock during the night. Such response would explain why the domesticated *ELF3hv* alleles, under NB, confer late flowering, however, under SB such genotypes accelerate plant development more significantly compared to genotypes that harbour wild alleles, as in the latter conditions the night is very short. However, such response it is visible only in a *ppd-H1* background as *Ppd-H1* confers early flowering under both conditions. In this context the sensitive allele *Ppd-H1* under long days seems to be less influenced by *ELF3*’s suppression. In addition, an independent pathway has been hypothesized where the allele behind *eam8* and the wild *ELF3hsp* allele induce early flowering independently from *Ppd-H1* (Boden et al., 2014; Faure et al., 2012; Zahn et al., 2023). This would explain why we observe early flowering phenotypes in the presence of *ELF3hsp* in a *ppd-H1* background.

In summary, this study has highlighted the importance of both *PPD-H1* and *ELF3* in the control of speed breeding in barley. The deployment of the HEB-25 population enabled alleles at these two loci to be fully explored in the context of speed breeding. Our findings will be particularly important for the deployment of SB in crop improvement programs that focus on the incorporation of new sources of genetic variation from wild relatives (Gramazio et al., 2021; Hao et al., 2020; Hernandez et al., 2020; Khan et al., 2023; Zhang et al., 2023). Data from this study predict that the deployment of this technology to accelerate generation time in breeding, will select against specific alleles in the genomic regions on chromosomes 1 and 2 where *PPD-H1* and *ELF3* are located, if late flowering genotypes will not be advanced to the next generation. Furthermore, a comparison of the allelic series at *PPD-H1* and *ELF3* (**Figure 3**) identified that domesticated alleles at these two loci, which are those that tend to be unintentionally selected against under rapid cycling conditions, also are likely to be associated with higher levels of plasticity that may guide and support their deployment in breeding programs designed to create climate resilient cultivars. Finally, genotypes that harbour wild alleles showed early flowering under both conditions, suggesting that, in such genotypes, speed breeding might not be as important to accelerate generation time as in domesticated genotypes.

## Acknowledgements

This project was funded by EASTBIO DTP and BBSRC to Nicola Rossi and by direct funding to Rajiv Sharma from SRUC. We thank Principal’s research group members Chin Jian Yang, Mike Smith, Ian Dawson, and David Marshall (SRUC) for helpful discussion throughout the work. Acknowledgements also go to Kalina Gorniak and the technicians (Julie Fortune, Grace Cuthill and Lachlan Jones) in the crops and soil department for helping with the experiments in the glasshouse.

## Supplementary Data

**Figure S1.** Principal component analysis for HEB-25 based on 32,955 SNP data.

**Figure S2.** Figure showing the frequency distribution of the 6 traits (a-f).

**Figure S3.** The plot displays a correlation matrix (from Table S2), with the six different traits represented along both the x-axis and the y-axis.

**Figure S4.** Manhattan plots from the six-traits using PPD-H1 and ELF3 as covariates.

**Figure S5.** The graph displays a distribution of a MAT SB across 162 entries, categorized into four distinct genotype groups. The x-axis represents percentiles. Each point is color-coded to represent one of the four genotype groups, allowing for a visual comparison of their distribution in the dataset.

**Data S1**. List of the BLUEs value of each of the 6 traits for the genotypes of the HEB 25 used in this study.

**Data S2** Significant (log10 p-val > 4) markers and their effect for the 6 GWAS for each of the trait analyzed

**Data S3a.** P-values resulted observed and permuted based data t-tests between genotype groups.

**Data S3b**. Summary statistics of the different genotype groups as in **Table 1**.

**Data S4**. List in decrescent order the genotypes and the genotype groups for the trait MAT SB.

**Data S5.** Significant (log10 p-val > 4) markers and their effect for the 6 GWAS, using ELF3 and PPD-H1 as covariates, for each of the trait analyzed.

**Table S1** Table showing the summary statistics of the 4 traits with mean, standard deviation, minimum, maximum and heritability estimates using Piepho’s method

**Table S2** Table showing Pearson correlation matrix between the 6 different traits.

## Notes

### Competing Interest Statement

The authors have declared no competing interest.

## References

Ahmar, S., Gill, R. A., Jung, K. H., Faheem, A., Qasim, M. U., Mubeen, M., & Zhou, W. (2020). Conventional and molecular techniques from simple breeding to speed breeding in crop plants: Recent advances and future outlook. International Journal of Molecular Sciences, 21(7), 1–24. 10.3390/ijms21072590

Arthur, J. M., Guthrie, J. D., & Newell, J. M. (1930). Some Effects of Artificial Climates on the Growth and Chemical Composition of Plants. American Journal of Botany, 17(5), 416. 10.2307/2435930

Bates, D., Mächler, M., Bolker, B., & Walker, S. (2015). Fitting Linear Mixed-Effects Models Using lme4. Journal of Statistical Software, 67(1). 10.18637/jss.v067.i01

Bayer, M. M., Rapazote-Flores, P., Ganal, M., Hedley, P. E., Macaulay, M., Plieske, J., Ramsay, L., Russell, J., Shaw, P. D., Thomas, W., & Waugh, R. (2017). Development and Evaluation of a Barley 50k iSelect SNP Array. Frontiers in Plant Science, 8. 10.3389/fpls.2017.01792

Bhatta, M., Sandro, P., Smith, M. R., Delaney, O., Voss-Fels, K. P., Gutierrez, L., & Hickey, L. T. (2021). Need for speed: manipulating plant growth to accelerate breeding cycles. Current Opinion in Plant Biology, 60, 101986. 10.1016/j.pbi.2020.101986

Boden, S. A., Weiss, D., Ross, J. J., Davies, N. W., Trevaskis, B., Chandler, P. M., & Swain, S. M. (2014). EARLY FLOWERING3 regulates flowering in spring barley by mediating gibberellin production and FLOWERING LOCUS T expression. Plant Cell, 26(4), 1557–1569. 10.1105/tpc.114.123794

Bohra, A., Chand Jha, U., Godwin, I. D., & Kumar Varshney, R. (2020). Genomic interventions for sustainable agriculture. Plant Biotechnology Journal, 18(12), 2388–2405. 10.1111/pbi.13472

Büttner, B., Draba, V., Pillen, K., Schweizer, G., & Maurer, A. (2020). Identification of QTLs conferring resistance to scald (Rhynchosporium commune) in the barley nested association mapping population HEB-25. BMC Genomics, 21(1), 837. 10.1186/s12864-020-07258-7

Caligari, P. D. S., Powell, W., & Jinks, J. L. (1987). A comparison of inbred lines derived by doubled haploidy and single seed descent in spring barley (Hordeum vulgare). Annals of Applied Biology, 111(3), 667–675. 10.1111/j.1744-7348.1987.tb02024.x

Campoli, C., Shtaya, M., Davis, S. J., & von Korff, M. (2012). Expression conservation within the circadian clock of a monocot: natural variation at barley Ppd-H1affects circadian expression of flowering time genes, but not clock orthologs. BMC Plant Biology, 12(1), 97. 10.1186/1471-2229-12-97

Cazzola, F., Bermejo, C. J., Guindon, M. F., & Cointry, E. (2020). Speed breeding in pea (Pisum sativum L.), an efficient and simple system to accelerate breeding programs. Euphytica, 216(11), 178. 10.1007/s10681-020-02715-6

Cha, J.-K., O’Connor, K., Alahmad, S., Lee, J.-H., Dinglasan, E., Park, H., Lee, S.-M., Hirsz, D., Kwon, S.-W., Kwon, Y., Kim, K.-M., Ko, J.-M., Hickey, L. T., Shin, D., & Dixon, L. E. (2022). Speed vernalization to accelerate generation advance in winter cereal crops. Molecular Plant, 15(8), 1300–1309. 10.1016/j.molp.2022.06.012

Cheng, J., Hill, C., Han, Y., He, T., Ye, X., Shabala, S., Guo, G., Zhou, M., Wang, K., & Li, C. (2023). New semi-dwarfing alleles with increased coleoptile length by gene editing of gibberellin 3-oxidase 1 using CRISPR-Cas9 in barley (Hordeum vulgare L.). Plant Biotechnology Journal, 21(4), 806–818. 10.1111/pbi.13998

Chiurugwi, T., Kemp, S., Powell, W., & Hickey, L. T. (2019). Speed breeding orphan crops. Theoretical and Applied Genetics, 132(3), 607–616. 10.1007/s00122-018-3202-7

Cooper, M., Tang, T., Gho, C., Hart, T., Hammer, G., & Messina, C. (2020). Integrating genetic gain and gap analysis to predict improvements in crop productivity. Crop Science, 60(2), 582–604. 10.1002/csc2.20109

Digel, B., Tavakol, E., Verderio, G., Tondelli, A., Xu, X., Cattivelli, L., Rossini, L., & von Korff, M. (2016). Photoperiod-H1 (Ppd-H1) Controls Leaf Size. Plant Physiology, 172(1), 405–415. 10.1104/pp.16.00977

Dixon, L. E., Knox, K., Kozma-Bognar, L., Southern, M. M., Pokhilko, A., & Millar, A. J. (2011). Temporal Repression of Core Circadian Genes Is Mediated through EARLY FLOWERING 3 in Arabidopsis. Current Biology, 21(2), 120–125. 10.1016/j.cub.2010.12.013

Ejaz, M., & von Korff, M. (2017). The Genetic Control of Reproductive Development under High Ambient Temperature. Plant Physiology, 173(1), 294–306. 10.1104/pp.16.01275

Endelman, J. B. (2011). Ridge Regression and Other Kernels for Genomic Selection with R Package rrBLUP. The Plant Genome, 4(3), 250–255. 10.3835/plantgenome2011.08.0024

Fang, Y., Wang, L., Sapey, E., Fu, S., Wu, T., Zeng, H., Sun, X., Qian, S., Khan, M. A. A., Yuan, S., Wu, C., Hou, W., Sun, S., & Han, T. (2021). Speed-Breeding System in Soybean: Integrating Off-Site Generation Advancement, Fresh Seeding, and Marker-Assisted Selection. Frontiers in Plant Science, 12. 10.3389/fpls.2021.717077

Faure, S., Turner, A. S., Gruszka, D., Christodoulou, V., Davis, S. J., Von Korff, M., & Laurie, D. A. (2012). Mutation at the circadian clock gene EARLY MATURITY 8 adapts domesticated barley (Hordeum vulgare) to short growing seasons. Proceedings of the National Academy of Sciences of the United States of America, 109(21), 8328–8333. 10.1073/pnas.1120496109

Fradgley, N., Gardner, K. A., Bentley, A. R., Howell, P., Mackay, I. J., Scott, M. F., Mott, R., & Cockram, J. (2023). Multi-trait ensemble genomic prediction and simulations of recurrent selection highlight importance of complex trait genetic architecture for long-term genetic gains in wheat. In Silico Plants, 5(1). 10.1093/insilicoplants/diad002

Watson, A., Ghosh, S., Williams, M.J. et al. Speed breeding is a powerful tool to accelerate crop research and breeding. Nature Plants 4, 23–29 (2018). 10.1038/s41477-017-0083-8

Gol, L., Haraldsson, E. B., & von Korff, M. (2021). Ppd-H1 integrates drought stress signals to control spike development and flowering time in barley. Journal of Experimental Botany, 72(1), 122–136. 10.1093/jxb/eraa261

Gosal, S. S., Pathak, D., Wani, S. H., Vij, S., & Pathak, M. (2020). Accelerated Breeding of Plants: Methods and Applications. In Accelerated Plant Breeding, Volume 1 (pp. 1–29). Springer International Publishing. 10.1007/978-3-030-41866-3_1

Gramazio, P., Prohens, J., Toppino, L., & Plazas, M. (2021). Editorial: Introgression Breeding in Cultivated Plants. Frontiers in Plant Science, 12. 10.3389/fpls.2021.764533

Hao, M., Zhang, L., Ning, S., Huang, L., Yuan, Z., Wu, B., Yan, Z., Dai, S., Jiang, B., Zheng, Y., & Liu, D. (2020). The Resurgence of Introgression Breeding, as Exemplified in Wheat Improvement. Frontiers in Plant Science, 11. 10.3389/fpls.2020.00252

He, T., Hill, C. B., Angessa, T. T., Zhang, X.-Q., Chen, K., Moody, D., Telfer, P., Westcott, S., & Li, C. (2019). Gene-set association and epistatic analyses reveal complex gene interaction networks affecting flowering time in a worldwide barley collection. Journal of Experimental Botany, 70(20), 5603–5616. 10.1093/jxb/erz332

Hernandez, J., Meints, B., & Hayes, P. (2020). Introgression Breeding in Barley: Perspectives and Case Studies. Frontiers in Plant Science, 11. 10.3389/fpls.2020.00761

Herrero, E., Kolmos, E., Bujdoso, N., Yuan, Y., Wang, M., Berns, M. C., Uhlworm, H., Coupland, G., Saini, R., Jaskolski, M., Webb, A., Gonçalves, J., & Davis, S. J. (2012). EARLY FLOWERING4 Recruitment of EARLY FLOWERING3 in the Nucleus Sustains the Arabidopsis Circadian Clock. The Plant Cell, 24(2), 428–443. 10.1105/tpc.111.093807

Hickey, J. M., Chiurugwi, T., Mackay, I., & Powell, W. (2017). Genomic prediction unifies animal and plant breeding programs to form platforms for biological discovery. Nature Genetics, 49(9), 1297– 1303. 10.1038/ng.3920

Hickey, L. T., Germán, S. E., Pereyra, S. A., Diaz, J. E., Ziems, L. A., Fowler, R. A., Platz, G. J., Franckowiak, J. D., & Dieters, M. J. (2017). Speed breeding for multiple disease resistance in barley. Euphytica, 213(3), 64. 10.1007/s10681-016-1803-2

Hickey, L. T., N. Hafeez, A., Robinson, H., Jackson, S. A., Leal-Bertioli, S. C. M., Tester, M., Gao, C., Godwin, I. D., Hayes, B. J., & Wulff, B. B. H. (2019). Breeding crops to feed 10 billion. Nature Biotechnology, 37(7), 744–754. 10.1038/s41587-019-0152-9

Hooghvorst, I., & Nogués, S. (2021). Chromosome doubling methods in doubled haploid and haploid inducer-mediated genome-editing systems in major crops. Plant Cell Reports, 40(2), 255–270. 10.1007/s00299-020-02605-0

Hu, H., Wang, P., Angessa, T. T., Zhang, X., Chalmers, K. J., Zhou, G., Hill, C. B., Jia, Y., Simpson, C., Fuller, J., Saxena, A., Al Shamaileh, H., Iqbal, M., Chapman, B., Kaur, P., Dudchenko, O., Aiden, E. L., … Li, C. (2023). Genomic signatures of barley breeding for environmental adaptation to the new continents. Plant Biotechnology Journal. 10.1111/pbi.14077

Inagaki, M. N., Varughese, G., Rajaram, S., van Ginkel, M., & Mujeeb-Kazi, A. (1998). Comparison of bread wheat lines selected by doubled haploid, single-seed descent and pedigree selection methods. Theoretical and Applied Genetics, 97(4), 550–556. 10.1007/s001220050930

Isidro-Sánchez, J., Akdemir, D., & Montilla-Bascón, G. (2017). Genome-Wide Association Analysis Using R (pp. 189–207). 10.1007/978-1-4939-6682-0_14

Khan, A. H., Min, L., Ma, Y., Zeeshan, M., Jin, S., & Zhang, X. (2023). High-temperature stress in crops: male sterility, yield loss and potential remedy approaches. Plant Biotechnology Journal, 21(4), 680–697. 10.1111/pbi.13946

Laitinen, R. A. E., & Nikoloski, Z. (2019). Genetic basis of plasticity in plants. Journal of Experimental Botany, 70(3), 795–804. 10.1093/jxb/ery404

Lanchashire, P. D., Bleiholder, H., Boom, T. Van Den, Langeluddeke, P., Stauss, R., Weber, E., & Witzemberger, A. (1991). A uniform decimal code for growth stages of crops and weeds. Annals of Applied Biology, 119(3), 561–601. 10.1111/j.1744-7348.1991.tb04895.x

Li, H., Rasheed, A., Hickey, L. T., & He, Z. (2018). Fast-Forwarding Genetic Gain. Trends in Plant Science, 23(3), 184–186. 10.1016/j.tplants.2018.01.007

Lundqvist, U. (2009). Eighty years of Scandinavian barley mutation genetics and breeding. Induced plant mutations in the genomics era. Food and Agriculture Organization of the United Nations, Rome, 39–43.

Maurer, A., Draba, V., Jiang, Y., Schnaithmann, F., Sharma, R., Schumann, E., Kilian, B., Reif, J. C., & Pillen, K. (2015). Modelling the genetic architecture of flowering time control in barley through nested association mapping. BMC Genomics, 16(1), 290. 10.1186/s12864-015-1459-7

Maurer, A., Draba, V., & Pillen, K. (2016). Genomic dissection of plant development and its impact on thousand grain weight in barley through nested association mapping. Journal of Experimental Botany, 67(8), 2507–2518. 10.1093/jxb/erw070

Maurer, et al. (2019-11-21): 50k Illumina Infinium iSelect SNP Array data for the wild barley NAM population HEB-25. DOI:10.5447/ipk/2019/20

Mehnaz, M., Dracatos, P., Pham, A., March, T., Maurer, A., Pillen, K., Forrest, K., Kulkarni, T., Pourkheirandish, M., Park, R. F., & Singh, D. (2021). Discovery and fine mapping of Rph28: a new gene conferring resistance to Puccinia hordei from wild barley. Theoretical and Applied Genetics, 134(7), 2167–2179. 10.1007/s00122-021-03814-1

Mobini, S., Khazaei, H., Warkentin, T. D., & Vandenberg, A. (2020). Shortening the generation cycle in faba bean (Vicia faba) by application of cytokinin and cold stress to assist speed breeding. Plant Breeding, 139(6), 1181–1189. 10.1111/pbr.12868

Müller, L. M., Mombaerts, L., Pankin, A., Davis, S. J., Webb, A. A. R., Goncalves, J., & Von Korff, M. (2020). Differential effects of day/night cues and the circadian clock on the barley transcriptome. Plant Physiology, 183(2), 765–779. 10.1104/pp.19.01411

Nusinow, D. A., Helfer, A., Hamilton, E. E., King, J. J., Imaizumi, T., Schultz, T. F., Farré, E. M., & Kay, S. A. (2011). The ELF4–ELF3–LUX complex links the circadian clock to diurnal control of hypocotyl growth. Nature, 475(7356), 398–402. 10.1038/nature10182

Ochagavía, H., Kiss, T., Karsai, I., Casas, A. M., & Igartua, E. (2022). Responses of Barley to High Ambient Temperature Are Modulated by Vernalization. Frontiers in Plant Science, 12. 10.3389/fpls.2021.776982

Pandey, S., Singh, A., Parida, S. K., & Prasad, M. (2022). Combining speed breeding with traditional and genomics-assisted breeding for crop improvement. Plant Breeding, 141(3), 301–313. 10.1111/pbr.13012

Piepho, H.-P., & Möhring, J. (2007). Computing Heritability and Selection Response From Unbalanced Plant Breeding Trials. Genetics, 177(3), 1881–1888. 10.1534/genetics.107.074229

Powell, W., Caligari, P. D. S., & Thomas, W. T. B. (1986). Comparison of Spring Barley Lines Produced by Single Seed Descent, Pedigree Inbreeding and Doubled Haploidy. Plant Breeding, 97(2), 138–146. 10.1111/j.1439-0523.1986.tb01045.x

Qaseem, M. F., Qureshi, R., & Shaheen, H. (2019). Effects of Pre-Anthesis Drought, Heat and Their Combination on the Growth, Yield and Physiology of diverse Wheat (Triticum aestivum L.) Genotypes Varying in Sensitivity to Heat and drought stress. Scientific Reports, 9(1), 6955. 10.1038/s41598-019-43477-z

Russell, J., Mascher, M., Dawson, I. K., Kyriakidis, S., Calixto, C., Freund, F., Bayer, M., Milne, I., Marshall-Griffiths, T., Heinen, S., Hofstad, A., Sharma, R., Himmelbach, A., Knauft, M., van Zonneveld, M., Brown, J. W. S., Schmid, K., … Waugh, R. (2016a). Exome sequencing of geographically diverse barley landraces and wild relatives gives insights into environmental adaptation. Nature Genetics, 48(9), 1024–1030. 10.1038/ng.3612

Russell, J., Mascher, M., Dawson, I. K., Kyriakidis, S., Calixto, C., Freund, F., Bayer, M., Milne, I., Marshall-Griffiths, T., Heinen, S., Hofstad, A., Sharma, R., Himmelbach, A., Knauft, M., van Zonneveld, M., Brown, J. W. S., Schmid, K., … Waugh, R. (2016b). Exome sequencing of geographically diverse barley landraces and wild relatives gives insights into environmental adaptation. Nature Genetics, 48(9), 1024–1030. 10.1038/ng.3612

Saade, S., Maurer, A., Shahid, M., Oakey, H., Schmöckel, S. M., Negrão, S., Pillen, K., & Tester, M. (2016). Yield-related salinity tolerance traits identified in a nested association mapping (NAM) population of wild barley. Scientific Reports, 6(1), 32586. 10.1038/srep32586

Samantara, K., Bohra, A., Mohapatra, S. R., Prihatini, R., Asibe, F., Singh, L., Reyes, V. P., Tiwari, A., Maurya, A. K., Croser, J. S., Wani, S. H., Siddique, K. H. M., & Varshney, R. K. (2022). Breeding More Crops in Less Time: A Perspective on Speed Breeding. Biology, 11(2), 275. 10.3390/biology11020275

Samineni, S., Sen, M., Sajja, S. B., & Gaur, P. M. (2020). Rapid generation advance (RGA) in chickpea to produce up to seven generations per year and enable speed breeding. The Crop Journal, 8(1), 164–169. 10.1016/j.cj.2019.08.003

Schilling, S., Melzer, R., Dowling, C. A., Shi, J., Muldoon, S., & McCabe, P. F. (2023). A protocol for rapid generation cycling (speed breeding) of hemp (Cannabis sativa) for research and agriculture. The Plant Journal, 113(3), 437–445. 10.1111/tpj.16051

Sharma, R., Draicchio, F., Bull, H., Herzig, P., Maurer, A., Pillen, K., Thomas, W. T. B., & Flavell, A. J. (2018). Genome-wide association of yield traits in a nested association mapping population of barley reveals new gene diversity for future breeding. Journal of Experimental Botany, 69(16), 3811–3822. 10.1093/jxb/ery178

Smith, P. (2013). Delivering food security without increasing pressure on land. Global Food Security, 2(1), 18–23. 10.1016/j.gfs.2012.11.008

Song, Y., Duan, X., Wang, P., Li, X., Yuan, X., Wang, Z., Wan, L., Yang, G., & Hong, D. (2022). Comprehensive speed breeding: a high-throughput and rapid generation system for long-day crops. Plant Biotechnology Journal, 20(1), 13–15. 10.1111/pbi.13726

Turner, A., Beales, J., Faure, S., Dunford, R. P., & Laurie, D. A. (2005). The Pseudo-Response Regulator Ppd-H1 Provides Adaptation to Photoperiod in Barley. Science, 310(5750), 1031–1034. 10.1126/science.1117619

W. W. Garner, and H. A. Allard (1927). Effect of Short Alternating Periods of Light and Darkness on Plant Growth. Science, Vf. LXVI N(1697). 10.1126/science.66.1697.40

Wanga, M. A., Shimelis, H., Mashilo, J., & Laing, M. D. (2021). Opportunities and challenges of speed breeding: A review. Plant Breeding, 140(2), 185–194. 10.1111/pbr.12909

Watson, A., Ghosh, S., Williams, M. J., Cuddy, W. S., Simmonds, J., Rey, M. D., Asyraf Md Hatta, M., Hinchliffe, A., Steed, A., Reynolds, D., Adamski, N. M., Breakspear, A., Korolev, A., Rayner, T., Dixon, L. E., Riaz, A., Martin, W., … Hickey, L. T. (2018). Speed breeding is a powerful tool to accelerate crop research and breeding. Nature Plants, 4(1), 23–29. 10.1038/s41477-017-0083-8

Wickham H (2016). ggplot2: Elegant Graphics for Data Analysis. Springer-Verlag New York. ISBN 978-3-319-24277-4, https://ggplot2.tidyverse.org.

Wiegmann, M., Maurer, A., Pham, A., March, T. J., Al-Abdallat, A., Thomas, W. T. B., Bull, H. J., Shahid, M., Eglinton, J., Baum, M., Flavell, A. J., Tester, M., & Pillen, K. (2019). Barley yield formation under abiotic stress depends on the interplay between flowering time genes and environmental cues. Scientific Reports, 9(1), 6397. 10.1038/s41598-019-42673-1

Yin, L., Zhang, H., Tang, Z., Xu, J., Yin, D., Zhang, Z., Yuan, X., Zhu, M., Zhao, S., Li, X., & Liu, X. (2021). rMVP: A Memory-efficient, Visualization-enhanced, and Parallel-accelerated Tool for Genome-wide Association Study. Genomics, Proteomics & Bioinformatics, 19(4), 619–628. 10.1016/j.gpb.2020.10.007

Zahn, T., Zhu, Z., Ritoff, N., Krapf, J., Junker, A., Altmann, T., Schmutzer, T., Tüting, C., Kastritis, P. L., Babben, S., Quint, M., Pillen, K., & Maurer, A. (2023). Novel exotic alleles of EARLY FLOWERING 3 determine plant development in barley. Journal of Experimental Botany. 10.1093/jxb/erad127

Zhang, W., Tan, C., Hu, H., Pan, R., Xiao, Y., Ouyang, K., Zhou, G., Jia, Y., Zhang, X., Hill, C. B., Wang, P., Chapman, B., Han, Y., Xu, L., Xu, Y., Angessa, T., Luo, H., … Li, C. (2023). Genome architecture and diverged selection shaping pattern of genomic differentiation in wild barley. Plant Biotechnology Journal, 21(1), 46–62. 10.1111/pbi.13917

Zhu, Z., Esche, F., Babben, S., Trenner, J., Serfling, A., Pillen, K., Maurer, A., & Quint, M. (2023). An exotic allele of barley EARLY FLOWERING 3 contributes to developmental plasticity at elevated temperatures. Journal of Experimental Botany, 74(9), 2912–2931. 10.1093/jxb/erac470

