## Supplementary Figures for "Investigating the genetic control of plant development under speed breeding conditions"

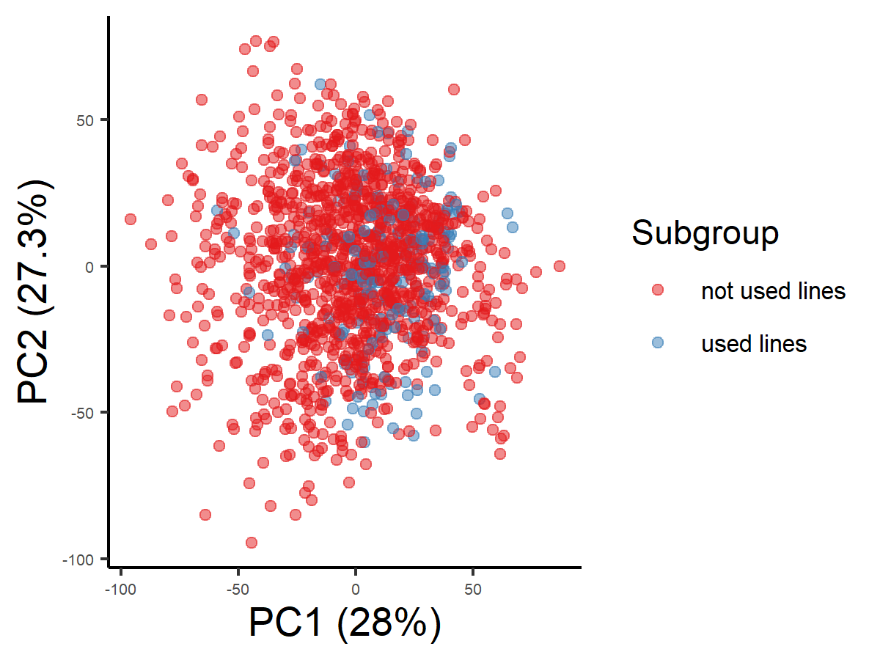
**Supplementary Figures**

**Figure S1**. Principal component analysis for HEB-25 based on 32,955 SNP data. Two principal components (PCs) capturing the largest amount of genetic variation are shown as x- and y-axiss, respectively. Percentages in brackets denote the variance explained by the respective PC. In red are the lines studies and in blue the entire HEB-25 population.

(b)

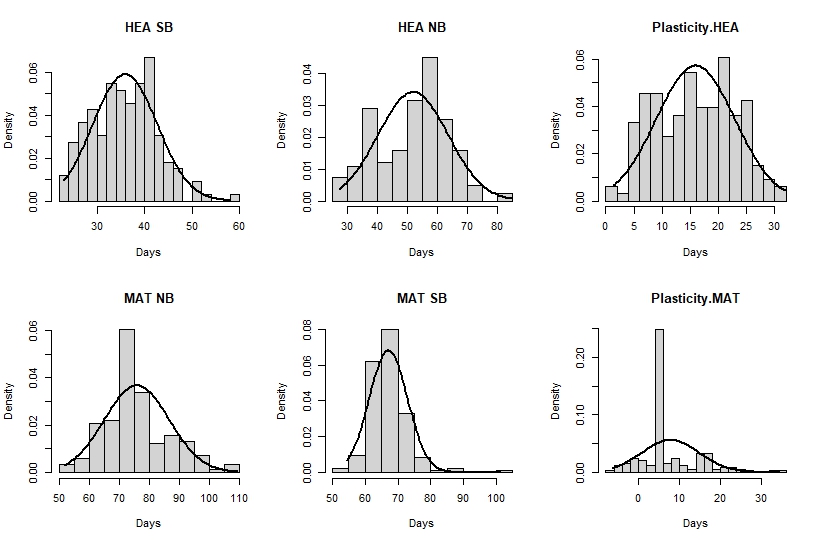

(d)

(e)

(f)

(c)

(a)

**Figure S2**. Figure showing the frequency distribution of the 6 traits (a-f).

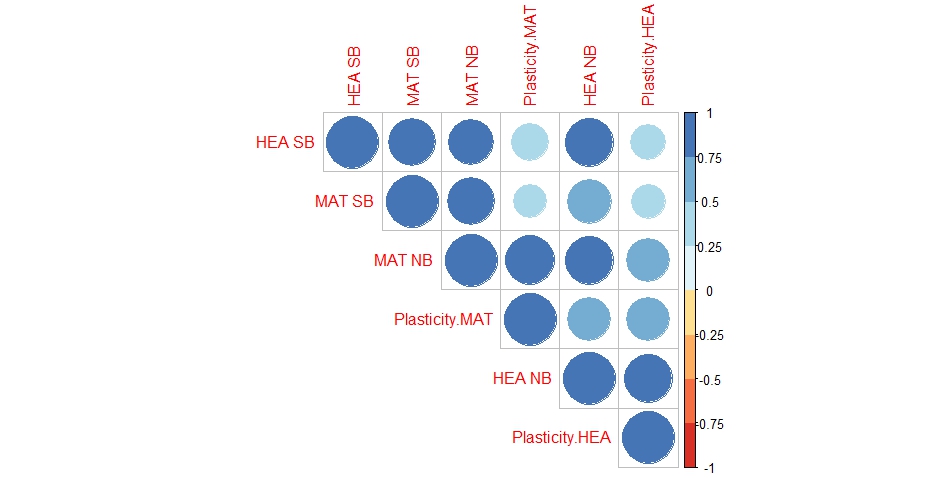

**Figure S3**. The plot displays a correlation matrix (from Table S2), with the six different traits represented along both the x-axis and the y-axis. Each cell in the matrix corresponds to the correlation coefficient between a pair of traits. The color of each cell represents the strength and direction of the correlation between the two traits. The color scale ranges from blue indicating a negative correlation, through white for no correlation, to a red indicating a positive correlation. The intensity of the color corresponds to the magnitude of the correlation coefficient, with darker shades indicating stronger correlations. The plot was created using the R -package “corrplot” (Wei T, Simko V 2021) in Rstudio version 4.2.2.

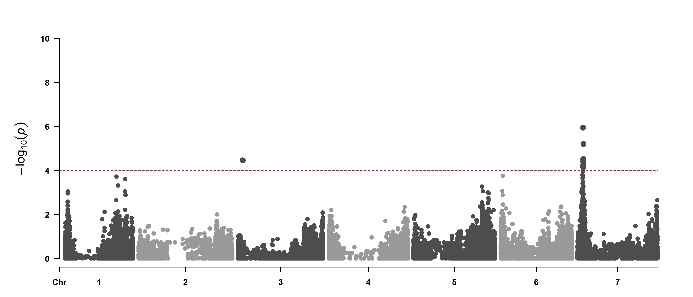

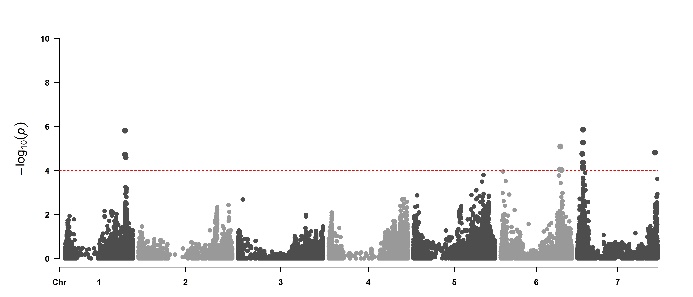

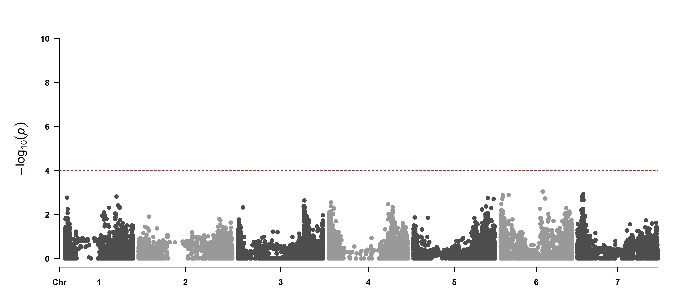

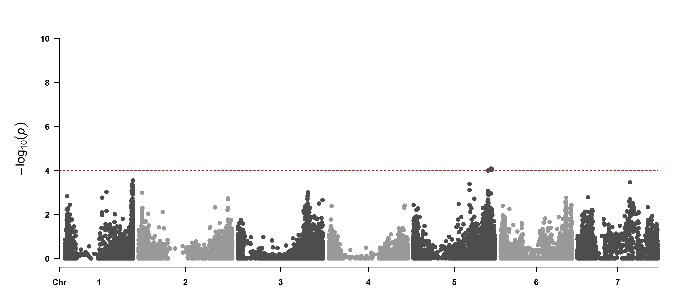

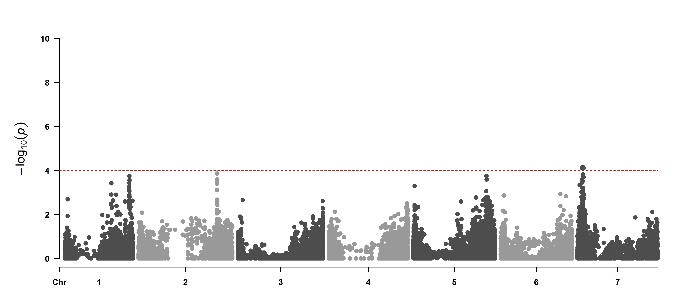

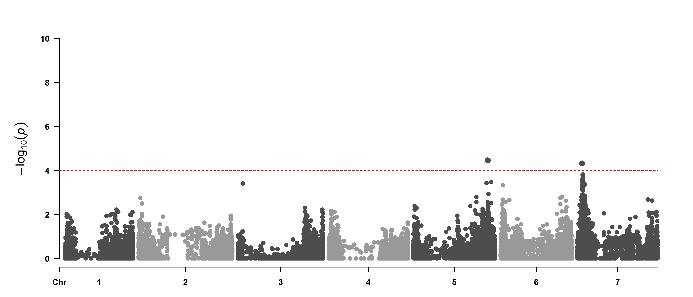

(d)

(a)

NB MAT

NB HEA

SB HEA

SB MAT

Plasticity.MAT

GA20ox/PPD-H2

CO2

FT1

Plasticity.HEA

(e)

(b)

(f)

(c)

**Figure S4**. Manhattan plots from the six-traits using PPD-H1 and ELF3 as covariates. Seven barley chromosomes are shown (1H-7H) horizontally and –log10(p-values) are displayed vertically by dotted line. Significant threshold dashed line set at –log10 (p-value) = 4.0. The coinciding flowering time candidate genes are shown in the rectangle boxes. Plots were created using the “CMplot” package (Yin et al., 2021) in R studio version 4.2.2. The details of the significant peaks and the markers underlying these peaks are provided in **Supplementary Table S5**.

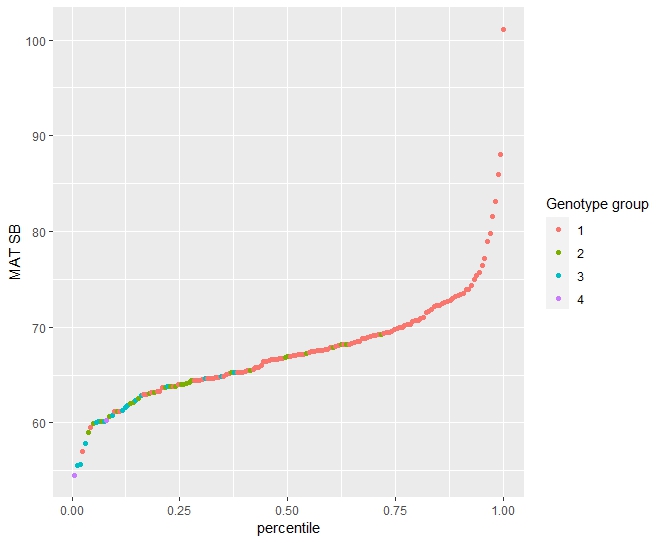

**Figure S5**. The graph displays a distribution of a MAT SB across 162 entries, categorized into four distinct genotype groups. The x-axis represents percentiles. Each point is color-coded to represent one of the four genotype groups, allowing for a visual comparison of their distribution in the dataset.
